# Exploiting Electrophysiological Measures of Semantic Processing for Auditory Attention Decoding

**DOI:** 10.1101/2020.04.17.046813

**Authors:** Karen Dijkstra, Peter Desain, Jason Farquhar

**Affiliations:** Radboud University, Donders Institute for Brain, Cognition and Behaviour, Nijmegen, the Netherlands; MindAffect, Nijmegen, the Netherlands

## Abstract

In Auditory Attention Decoding, a user’s electrophysiological brain responses to certain features of speech are modelled and subsequently used to distinguish attended from unattended speech in multi-speaker contexts. Such approaches are frequently based on acoustic features of speech, such as the auditory envelope. A recent paper shows that the brain’s response to a semantic description (i.e., semantic dissimilarity) of narrative speech can also be modelled using such an approach. Here we use the (publicly available) data accompanying that study, in order to investigate whether combining this semantic dissimilarity feature with an auditory envelope approach improves decoding performance over using the envelope alone. We analyse data from their ‘Cocktail Party’ experiment in which 33 subjects attended to one of two simultaneously presented audiobook narrations, for 30 1-minute fragments. We find that the addition of the dissimilarity feature to an envelope-based approach significantly increases accuracy, though the increase is marginal (85.4% to 86.6%). However, we subsequently show that this dissimilarity feature, in which the degree of dissimilarity of the current word with regard to the previous context is tagged to the onsets of each content word, can be replaced with a binary content-word-onset feature, without significantly affecting the results (i.e., modelled responses or accuracy), putting in question the added value of the dissimilarity information for the approach introduced in this recent paper.

## Introduction

Auditory attention enables us to attend to a single auditory source while ‘tuning out’ other sources^1, 2^. This ability allows us to function in the so-called ‘Cocktail Party’ situation, where multiple conversations are held simultaneously, yet we are able to perceive and understand the speech of a single speaker of interest. Recent developments in the neuroscience of understanding of speech have shown that in such a scenario certain features of the attended speech are represented more strongly in electrophysiological brain activity, than features of the unattended speech^3–6^.

These differences in neural representations of attended and unattended speech have subsequently been shown to allow us to decode from brain activity which speaker someone is paying attention to: auditory attention decoding^7–11^. This provides a proof of concept of the neurosciencific phenomenon, but also opens the path to develop neurotechnology that exploits this decoding. Such attentional decoding, in combination with technologically driven source separation of speech sources, could for instance lead to the development of hearing aids that selectively amplify the speaker of interest based on the user’s attention^12^.

Auditory attention decoding approaches take advantage of features of the speech signal that are known to be selectively enhanced for attended over unattended speech. Examples of such speech features are for instance the changes in sound intensity over time (the speech envelope)^7–9, 11^, or the changes in intensity over time across frequency bands (the speech spectogram)^12, 13^. Generally, a (regularized) regression is used to learn a mapping from the subject’s electrophysiological data (e.g., EEG), to the chosen speech signal features based on data for which the attended speech is known.

This mapping from electrophysiological data to speech signal features can then be applied to EEG data for which the attended source is unknown, to produce an estimate of the signal features of the attended speech fragment. The estimated attended speech signal features can be compared to features extracted from the candidate speech fragments that the subject was exposed to when the EEG was recorded. The candidate whose stimulus feature matches most closely (e.g., is more correlated) to the estimated feature is predicted to belong to the attended speech fragment. In a real-life scenario those candidate speech fragments will have to be obtained by performing source separation on the mixed audio, but in experimental settings, the unmixed speech fragments are often available.

The above presented approach, in which the brain’s response is used to predict the original stimulus features, is referred to as backward modelling. The same decoding approach can be performed using a forward model, in which the stimulus feature is mapped to the EEG. While forward models are more straightforward to interpret^14^, Wong et al., demonstrate how for (regularized) attention decoding based on the speech envelope the backward modelling approach is superior, achieving both a better fit (i.e., higher correlation) with the attended speech, and a higher decoding accuracy^11^.

A recent publication describes how the brain response representing the semantic processing of presented speech can be modelled analogously by building a stimulus feature vector that tags the onset of each content word with its dissimilarity to the preceding context^15^. Content words, here, broadly refer to the words in the sentence that convey the meaning, as opposed to words with a primarily grammatical function (i.e., function words). This brain response modelled in response to these dissimilarity values presents as a negative going wave in the 300-600 ms time range, and shares characteristics with the N400, an event related potential associated with the semantic processing of stimuli^16^. In the paper the authors train (amongst others), (forward) models on the dissimilarity values of attended speech and models on unattended speech. When these models are subsequently used to for estimation, the attended speech model provides significantly higher correlations between the predicted and true EEG than the unattended speech model.

However, the question remains to which degree this dissimilarity feature can be used for auditory attention decoding, and if so, whether it provides additional value over currently used features (e.g., the speech envelope). To answer this question we train models for both a semantic dissimilarity feature and a speech envelope feature to identify attended speech. We investigate whether the dissimilarity model alone can achieve better than chance accuracy, and whether combining the dissimilarity and speech envelope models works better than using the speech envelope model alone.

To answer these questions we make use of the data from the Broderick et al. study that has been made available online^17^. Specifically, we use data from the ‘Cocktail Party’ dataset, in which 33 subjects attended to one of two simultaneously presented audiobook narrations, for 30 one minute fragments.

## Results

To determine whether the addition of a semantic dissimilarity feature can improve auditory attention decoding, we compute the accuracy with which the attended stimulus can be predicted based on the semantic feature, the envelope feature, and a combination of both. We illustrate how we obtain such accuracy for a given stimulus feature in Figure 1. For each subject, we first train a (regularized) linear regression to predict the stimulus feature from their (z-scored) EEG, for each time-point. At test time, the trained weights can then be applied to new EEG data to obtain an estimation of the respective stimulus feature that corresponds to the (unknown) attended speech. By correlating this estimation to the stimulus feature extracted from the two candidate speech fragments, a prediction for the attended source can be made.

**Figure 1.**
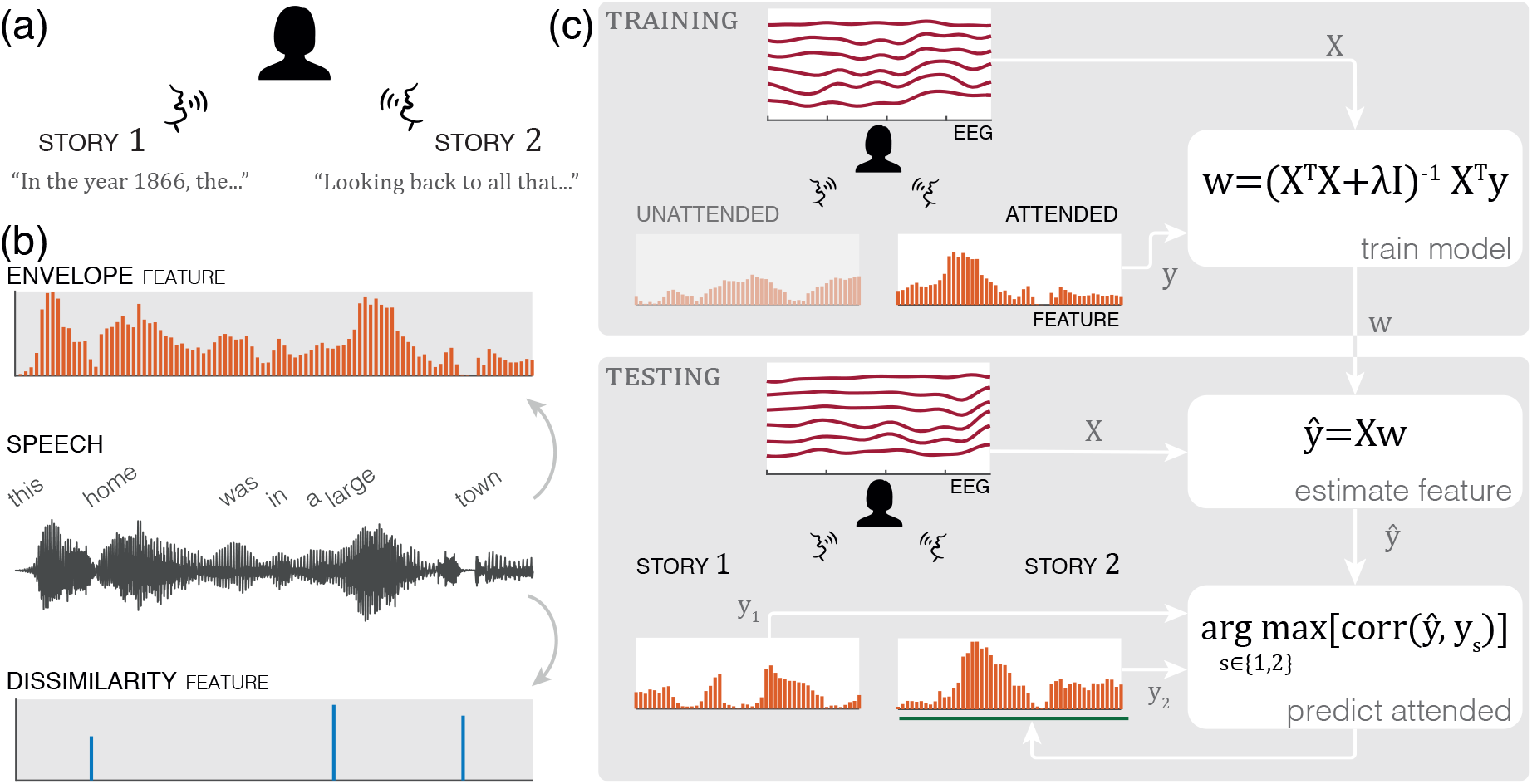
(**a**) Auditory Attention Paradigm. (**b**) Feature extraction. From the presented speech we extract the features of interest: the auditory envelope of the speech signal, and the semantic dissimilarity of content words tagged to their word onset. The non-content words, generally function words, are ignored. (**c**) Training: A given stimulus feature (here, the envelope), together with the EEG, is used to train the weights of a regression model, using a moving window of time lags (0-800ms). Testing: we have EEG data and two presented speech stimuli, but which of the two was attended is unknown. By applying the model (i.e., trained weights *w*) to this EEG data, an estimation of the attended stimulus feature ŷ is obtained. These are correlated to the relevant stimulus feature of the two candidate stimuli. The speech stimulus corresponding to the highest correlation is predicted to have been attended.

### Decoding results

Results for the identification of the attended speech can be found in Figure 2. Here we show the accuracy for the model trained on the semantic dissimilarity feature, a model trained on the speech envelope feature and a combined model. The accuracy of the combined model (env+sem) is obtained by summing the correlations of the two individual models, prior to prediction of the attended stimulus. This could be considered a naive approach, but we note that a (crossvalidated) classifier trained to weight the respective feature correlations did not improve decoding performance (p = 0.458, one sided, *α* = 0.05, randomization test, see *Methods*).

**Figure 2.**
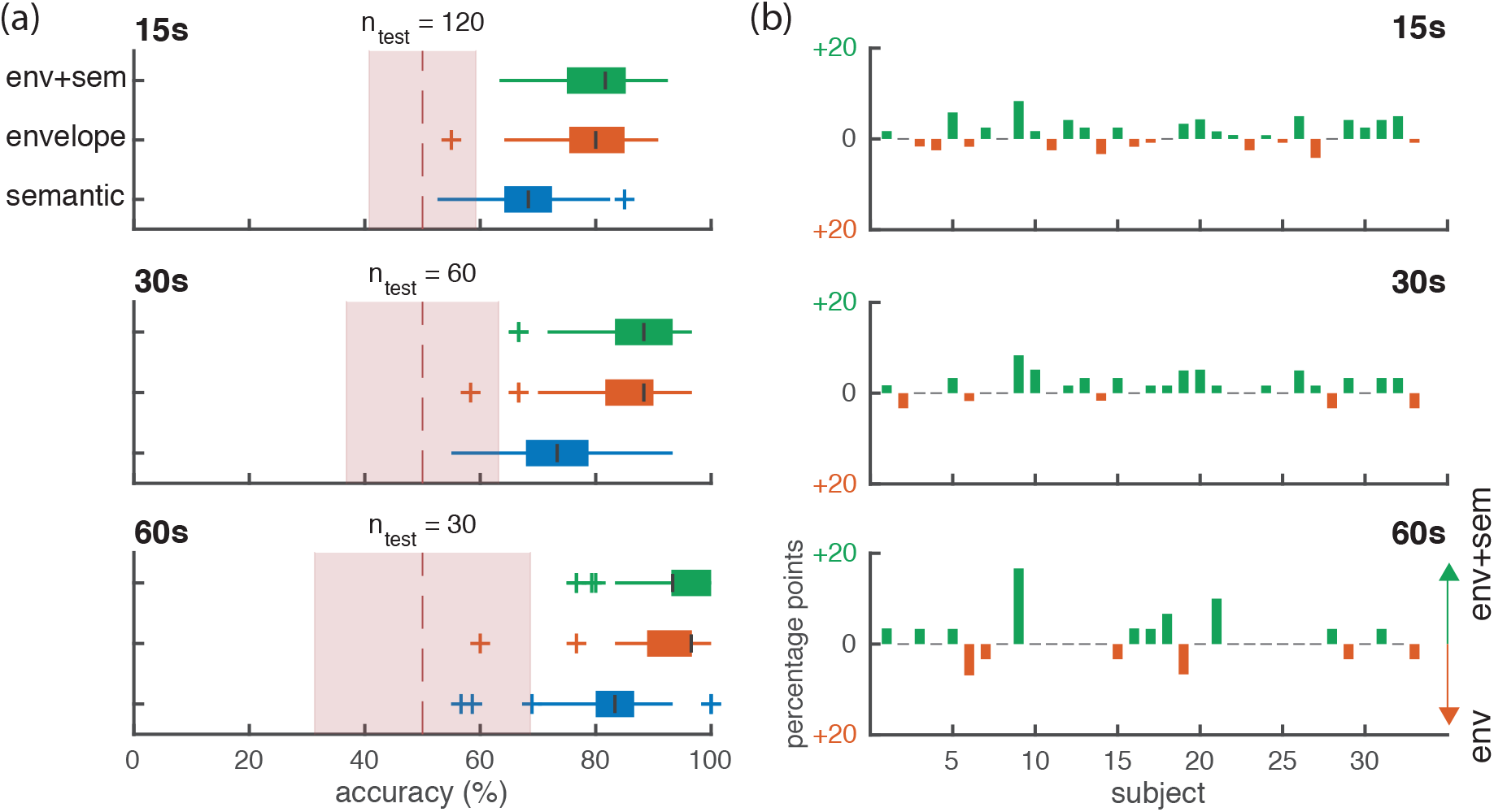
(**a**) accuracy of identifying the correct stimulus as attended, based on a model trained on either the semantic dissimilarity, the speech envelope or the combination of both (summed correlations), across all subjects, for three different lengths of the test segments (15, 30 and 60 s). Segments were created by taking non-overlapping slices of the full (60s) trials, resulting in a varying number of segments across durations (n_test_). The red shaded area depicts the 95% binomial confidence interval of chance accuracy (50%), based on the number of test segments considered (n_test_). (**b**) differences in accuracy between the speech envelope and combined (env+sem) model, per subject. An upward bar reflects a higher accuracy for the combined model, a downward bar a higher accuracy for the envelope model, in percentage points.

These accuracies are obtained using a nested crossvalidation of the available trials. In each test fold, the full trials (60 s) were evaluated at three segment lengths: 15, 30 and 60 seconds. Segments were created by taking non-overlapping slices of the full trials, resulting in a varying number of segments across durations. In the figure, we also plot a 95% binomial confidence interval for the chance level (50%), taking into account the respective number of observations (test segments) per subject. All three models achieve a median accuracy outside of the chance accuracy confidence interval. Note that five subjects were missing one out of 30 full trials, so the n_test_ variable represents the *median* number of trials per subject, and that the confidence interval is thus not always suitable for drawing inference with regard to a single subject.

To determine whether there is an increase in accuracy when using the combined model over the envelope model, we run a randomization test (1000 randomizations; see *Methods*). Comparing the observed difference we obtained between our models (*μ_comb_* = 86.61% – *μ_env_* = 85.44% = 1.17 percentage points), to the null-distribution of the randomizations, we find a significant difference (p = 0.001, one-sided, *α* = 0.05).

### Modelled brain responses

The weights from the trained models represent the filter that maps the EEG signal to the stimulus feature in question. The weights from a backward model are not directly interpretable in terms of the brain activity in response to that stimulus feature, but Haufe et al. provide an approach for transforming these weights to an interpretable activation pattern^14^. These are then comparable to the Temporal Response Functions (TRFs) obtained from forward models (e.g., those reported by Broderick et al.^15^). We plot these TRFs for the semantic dissimilarity and speech envelope in Figure 3. In Figure 3a and 3c we plot the time course together with a 95% bootstrapped confidence interval, for channels Fz and Pz respectively. In Figure 3b and 3d we plot the timecourse for all channels, using a colormap to reflect the TRF amplitude.

**Figure 3.**
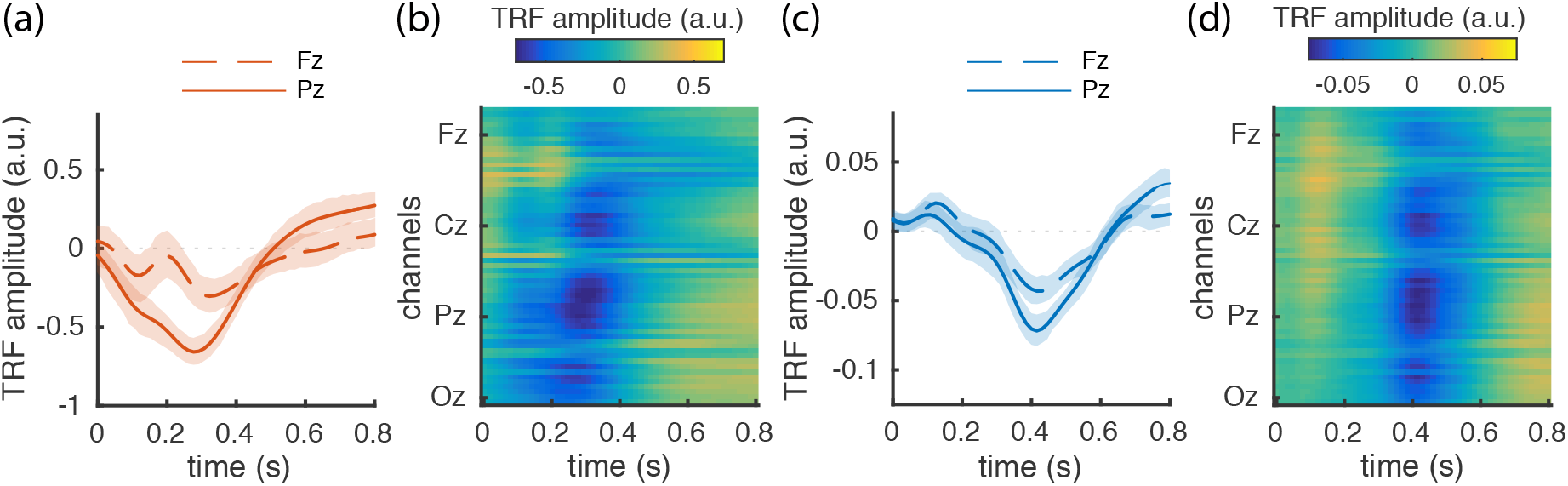
The weights of the backward models, transformed into (forward) Temporal Response Functions (TRFs). (**a** & **c**) The TRFs for two channels are plotted, for the speech envelope (**a**), and semantic dissimilarity model (**c**). The shaded regions represent the bootstrapped 95% confidence interval of the respective means (across subjects). (**b** & **d**) The TRFs across all channels, for the envelope (**b**) and dissimilarity model (**d**), ordered from anterior to posterior (top to bottom, respectively).

For the model based on the speech envelope there is an initial (100-200ms) positivity for fronto-central channels, and a broad negativity in the 200-400 ms, strongest in the centro-parietal channels. Finally, there is a a positivity across the parietal occipital channels in the 500-800 ms range. The TRF in response to the semantic dissimilarity, has a broader fronto-central positivity (100-200 ms), followed by a negativity in the 300-600 ms range). Note that amplitude of these TRFs is dependent on both the (z-scored) EEG and the feature values, so the two features’ TRF amplitudes are not comparable.

### Contribution of word onset

The semantic dissimilarity feature consists of the values of the dissimilarity of a given content word with respect to the preceding context, tagged to the onset of the word in question. The results reported above show that there is a brain response to this feature (Figure 3) and that combining this feature with the audio-envelope results in marginally improved attended speech identification (Figure 2).

By referring to it as a dissimilarity feature, it is suggested that it is the context-based dissimilarity that is driving the brain response. However, this feature implicitly encodes other information about the (semantic) content of the sentence, such as the onsets of the words and in particular, the onsets of the content words *only*. In the neuroscientific literature the most likely ERP corresponding to the brain response to this semantic feature is the N400. Previous work in this field suggests that open-class (i.e., content words) generally elicit a degree of N400 activity even in a congruent context, and that these open-class words result in a more negative N400 than closed-class words (i.e., function words) do^18^. Therefore, as a post-hoc question not addressed by Broderick et al., we investigate the relative contribution of these aspects in the resulting TRFs^15^.

First, we created an additional feature that tags the word onset of content words with a constant value. For comparison purposes we use the mean value of the dissimilarity feature values (see Fig. 4a). We train backward models for both the dissimilarity and the word onset feature to determine the relative contribution, and plot the resulting TRFs in Figure 4b.

**Figure 4.**
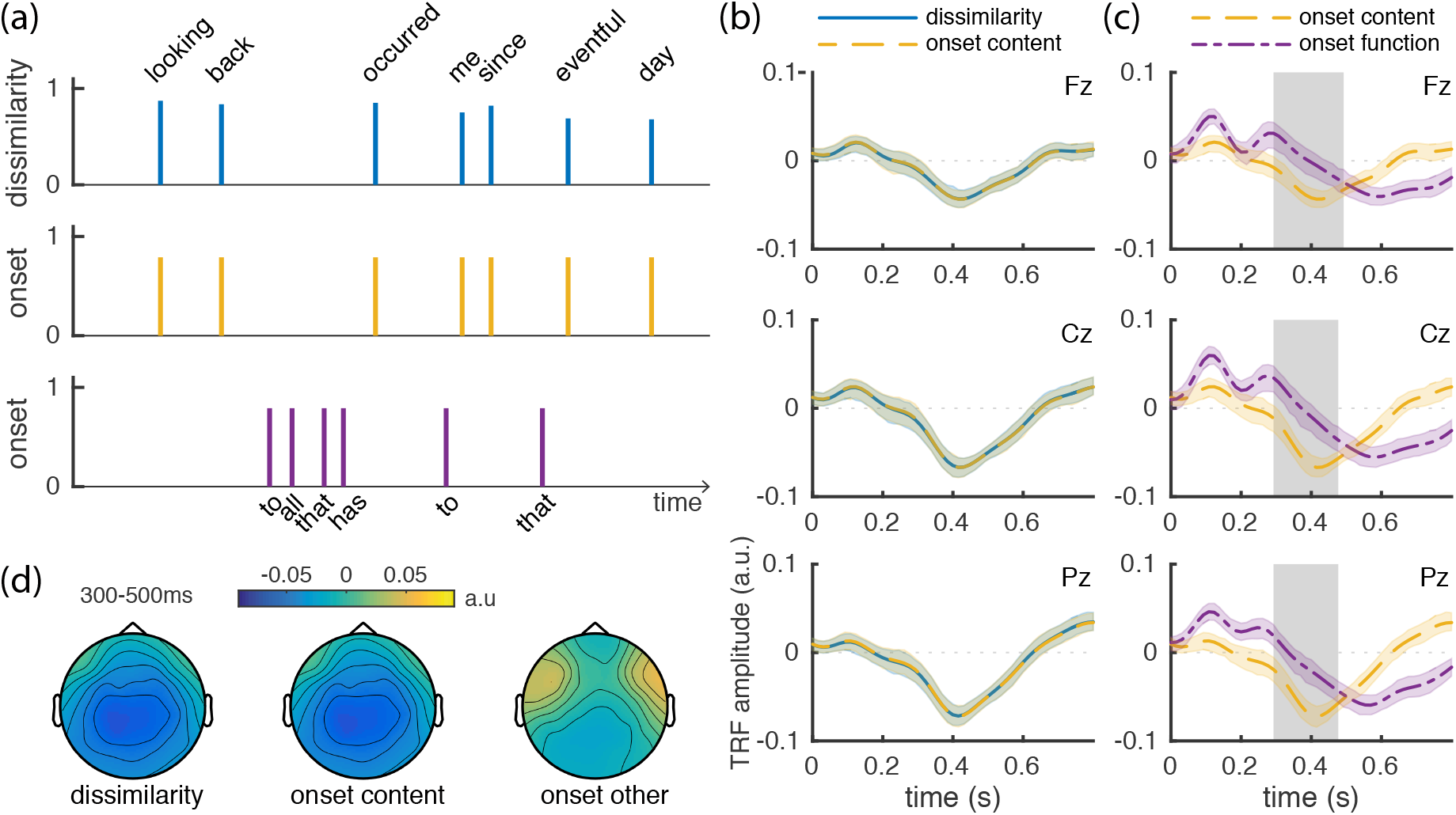
Comparison between the semantic dissimilarity model and models based on word onset only. (**a**) Features based on dissimilarity, the onset of content words, and the onset of function words, from top to bottom, respectively. (**b**) TRFs for the dissimilarity and the content word onset models. (**c**) TRFs for the onset models of the content words and function words. Shading represents the 95% bootstrapped confidence interval of the mean (across subjects). Boxes denote significant clusters of a non-parametric cluster-based permutation test obtained from comparing the respective TRFs. (**d**) Topographical plots for the dissimilarity, content onset and non-content onset, in the 300-500ms time range

We use a non-parametric cluster-based test to determine whether there is any difference between the TRF of the content word onset feature, and the content word dissimilarity feature in the 300-500ms^19^. This is based on the specific hypothesis that if the dissimilarity feature drives the semantic response, as indexed by the N400^16^, it would have a stronger negativity in the 300-500 ms range. We find no significant negative clusters (smallest p = 0.132, one-sided, alpha = 0.05). Furthermore, the model based on the content word onsets achieves the same mean accuracy (75.13%) as the dissimilarity based model (75.03%).

Next, we perform a post-hoc comparison to a feature indexing the onsets of the *non*-content words, with the aim to determine whether the word onset TRFs represent the activity to any arbitrary word, or content words in particular. If the latter were the case we would expect that the content words should show a more negative response in the 300-500ms range. The TRFs in response to the onset of content and the onset of non-content (i.e., function) words, can be found in Figure 4c. We test the difference between these in the same 300-500 ms window and find a significant negative cluster (p = 0.001, one-sided, alpha = 0.05), representing the lower amplitude in this time range for the content words.

We plot topographies of this 300-500 ms range in Figure 4d. Here we see a broad similarity between the dissimilarity and ‘onset content’ activity, and a reduction in centro-parietal negativity, and a frontal positivity for the function word onsets.

## Discussion

The aim of this study was to determine the potential benefit of the addition of a semantic dissimilarity feature for the purposes of auditory attention decoding, over the use of the speech envelope alone. We obtain three main results: 1) the dissimilarity feature on its own is sufficient to achieve better-than-chance accuracy, 2) there is a significant (but marginal) benefit from combining models in comparison to the envelope-only model and 3) when attempting to quantify the contribution of the word onsets to the TRFs of the dissimilarity feature, we find that these are highly similar, suggesting that it is not primarily the dissimilarity information that is driving this brain response, but rather the onset information of these content words.

On average, the combination of the dissimilarity and envelope model increases classifier accuracy by 1.2 percentage points (85.44% to 86.61%). A larger increase in accuracy could have been expected from combining models if the two features were completely independent in their effect on brain activity. This is unlikely to fully be the case (e.g., if the subject’s attention lapses for an audio fragment this would affect the brain response, and hence model predictions, for both features). Furthermore, the features can also be (partially or fully) dependent, in that they both encode the same information from the speech signal that the brain is responding to. The fact that the dissimilarity model’s TRFs are so similar to the TRFs of the word onset model may be an indication that the latter is the case, as it is more plausible that the speech envelope implicitly encodes the word onsets, than that it encodes contextual dissimilarity.

In fact, we can re-use the same regression approach to determine if this is the case: by fitting a model that predicts the content-word onsets on the basis of the speech envelope (rather than EEG). We use a crossvalidation approach to generate predictions and correlate these to the true onset vector, finding higher correlations for the matching (true) onsets than onsets of the other text (0.118 vs 0.002, respectively). While a seemingly weak correlation, this in fact a stronger correlation than obtained from using EEG as a predictor (0.040, for the best subject). The envelope feature, thus, indeed encodes the word onsets of content words to some degree, and to such a degree that it may explain the limited increase in performance from adding the semantic model/dissimilarity information.

For this study we used the existing Cocktail Party dataset from Broderick et al.^15, 17^. However, this experiment’s design has an important limitation when used for within-subject decoding of auditory attention: The experiment fixes the attended speaker, story and presentation side per subject, varying it only *between* subjects. This means that the trained models could have learned to exploit characteristics of the stimulus features that depend on speaker, story, or side, rather than attention. We do not know to which degree this has affected our reported decoding accuracies, but the median decoding accuracies for our speech envelope model are, if anything, somewhat lower than the comparable work of Wong et al.^11^, who do vary these factors within-subject. Furthermore, while the accuracies may not generalize to those from a realistic cocktail party scenario, we see no plausible mechanism for this to confound the conclusion with regard to the relative contribution of the semantic dissimilarity feature to the brain response TRFs.

The high similarity between the content word onset feature and the dissimilarity feature puts into question to what degree these TRFs indeed represent “electrophysiological correlates of semantic dissimilarity”, described as such by Broderick et al.^15^ The authors do provide a figure in their supplementary materials that shows a graded response to increasing values of dissimilarity, derived by binning the dissimilarities and re-obtaining a TRF per bin. This may suggest that the contribution of the dissimilarity is not zero, however, they provide no statistical analysis, and our results suggest this contribution, if anything, is small.

On the other hand, the fact that the TRFs to content and non-content words *are* significantly different, suggests that the TRFs also do not represent purely a word onset response. Furthermore, the TRFs of these content word or dissimilarity features have a similar waveform to that of the N400 that is known to be sensitive to semantic manipulations^16^. On the basis of the N400 literature, we could also predict a difference between the responses to the content and the non-content words, as a smaller negativity of the N400 has been observed for closed-class words (i.e., function words)^18^. Other ERP differences have been observed between the two word classes (open vs. closed), see e.g., Munte et al^20^. An example is the N400-700 that may correspond to the late (>400ms) negativity for function words seen in Fig. 4. Note however that the N400 is not only sensitive to semantic manipulations of context, but also to inherent word properties, such as word frequency or length that can correlate with word class. Overall, we think there is no reason to question that the Broderick et al. TRFs reflect the semantic processing of the (attended) natural speech, at least in as much as the N400 is an index thereof, even if the contribution of the dissimilarity measure itself is minimal.

Furthermore, the fact that we find the contribution of the dissimilarity information to the decoding of auditory attention here to be small-to-non-existent, does not mean that using features to describe variations in semantic context of speech cannot be successfully exploited for these purposes. As Broderick et al. note^15^, there are certainly more sophisticated measures of dissimilarity possible, using e.g., different definitions of ‘context’ (the words against which the dissimilarity is calculated), or different ways of encoding this dissimilarity in the features. Furthermore, the type of word-embeddings (i.e., word2vec) exploited for this dissimilarity measure are a relatively recent advancement in computational linguistics, and further advancements in that field could lead to better descriptors of semantic context to regress against.

Interestingly, even the function word onset feature can be exploited to identify the attended speaker from the unattended (mean accuracy 72.9%). This suggests that there are other linguistic descriptions of speech that could potentially be exploited for attention decoding. An exploration of such features and whether they can further increase auditory attention decoding accuracy would be a promising area for for further research.

In conclusion, with the current method of extracting a dissimilarity measurement from narrative speech, the semantic dissimilarity feature only marginally improves auditory attention decoding over using the speech envelopes alone. More sophisticated methods for extracting a continuous measure of the semantic content of (narrative) speech, may yet lead to improvements in the future, as may other linguistic descriptive features. Importantly, the current dissimilarity information does not contribute much, if anything, beyond the content word onset information, which means that anyone interested in using the effect outlined by Broderick et al. as a measure of semantic processing, may be served just as well by only determining the (content) word onsets.

## Methods

### Experimental data

Data were obtained from the Broderick et al. public dataset^17^. We analyse data from the ‘Cocktail Party’ dataset in which subjects attended to one of two simultaneous narrations of two separate audiobooks. The experimental data consists of 1) the 128 channel EEG data, recorded with the Biosemi Active 2 system, 2) the envelopes extracted from audio signals of the audiobook fragments that were used as stimuli and 3) the wordforms for the spoken text together with word onset and offset information. In the experiment, 33 subjects attended to one of two simultaneously presented audiobook narrations, for 30 one minute fragments. The two audiobooks, “Twenty thousand leagues under the sea” and “Journey to the Center of the Earth” by Jules Verne, were each narrated by different (male) speakers and presented on the left and right side respectively. 17 subjects attended the left, and 16 the right side.

According to the authors: “All procedures were undertaken in accordance with the Declaration of Helsinki and were approved by the Ethics Committees of the School of Psychology at Trinity College Dublin, and the Health Sciences Faculty at Trinity College Dublin”^15^. This included obtaining informed consent from all participants^21^. For further details on the experiment, please refer to Broderick et al.^15^, or the study by Power et al.^21^, for which the data was originally collected.

### EEG Preprocessing

The available EEG data consisted of 128 channels, sampled at 128 Hz, sliced by trial for each subject. Two additional mastoid channels were recorded to serve as a reference. We preprocessed this data by first highpass filtering at 0.5 Hz (4th order butterworth filter) and re-referencing the data to the mastoids electrodes. We chose a 0.5 Hz filter, rather than the 1 Hz chosen by Broderick et al., as the N400 has been shown to be sensitive to higher high-pass filters, but do not go as low as the 0.1 Hz suggested by Tanner et al., as we judged on the basis of a visual inspection that this resulted in too many low frequency artefacts^22^. We then regressed out any eye movement related activity by selecting the 5 frontal channels (C29, C30, C17, C16 and C8; roughly corresponding to Fp1, AF7, Fpz, Fp2 and AF8) as a proxy for EOG channels^23^. These channels were subsequently removed from the data. Next, we lowpass filtered the data at 8 Hz (4th order butterworth). A filter of 8 Hz has previously been used for the modelling of a brain response to both the semantic feature^15^, and a speech envelope feature^7^. A standard deviation based approach was used to identify bad channels (>3.5 deviations above the mean channel std), and any bad channels were then replaced by their interpolated equivalents using a spherical spline interpolation^24^. Note, for some subjects, a number of channels had an unusually small amplitude for the duration of the trials, prior to re-referencing; we treated these channels as bad channels in our analysis. Finally, we z-score the EEG (for each trial independently), to follow the procedure used by Broderick et al.

For computational efficiency, the data was then down sampled to 64Hz, and reduced from 123 (128-5) channels to a subset of channels most closely matching a biosemi 64 channel layout. The 5 frontal channels used to remove eye movement related activity were omitted, for a total of 59 channels. Given the 8 Hz lowpass filter, the data can be reduced to a 64Hz sampling frequency without loss of information, and a 64 channel layout has e.g., been shown sufficient for auditory attention decoding based on speech envelopes^11^.

### Speech feature extraction

The available stimulus data consisted of speech envelopes, sampled at 128Hz, together with a transcription of the content words in each speech fragment and their respective onset time. These speech envelopes were extracted by Broderick et al. from the raw audio signals using the (absolute of) Hilbert transform and time-locked to the EEG. We downsampled these speech envelopes to 64 Hz to match the sampling rate of the EEG. To create the semantic dissimilarity vector on the basis of the speech, we computed the semantic dissimilarity for each content word in the speech fragments as described in Broderick et al.^15^ For determining which were the content words, we followed the categorization of Broderick et al., that came pre-specified in the stimulus files in the public dataset. We obtained word vectors from the pre-trained word2vec model^25^, also publicly available, that Broderick et al. reported to use. This word2vec model defines 400-dimensional word vectors that can be used to calculate (cosine) similarity between the vectors of any arbitrary pair of words or phrases. For more details, refer to Baroni et al.^26^

The semantic similarity for a given word in the speech stimuli was calculated by taking the word vector of the respective word and correlating it to a phrase vector representing the previous context. This phrase vector was constructed by averaging the word-vectors of the previous words in the *current* sentence, or the average of the word-vectors in the *previous* sentence when the current word was the first content word in the sentence. The *dis*similarity was then computed by subtracting this similarity from one, resulting in a value between 0 and 2, with higher values indicating higher dissimilarity to the previous context. The semantic dissimilarity feature was created by tagging the onset of each content word with this dissimilarity value.

For creation of our word onset features (‘content’ or ‘function’), we use the words listed as content words by Broderick et al. for the content word feature. For the function words we then take all words not classed as content by that definition, and use a function-word list to remove proper nouns and other words of unknown status. These latter words would presumably have been excluded by Broderick et al. from the content word list when there was no word2vec vector available, even though these would normally be classed as open-class or content words.

### Decoding

We determine the decoding accuracy for a given subject and a given stimulus feature, with a nested crossvalidation (10-by-3 folds). For a given trial’s EEG and (attended) stimulus feature we train a backward model using the mTRF toolbox^13^(version 1.5). We train backward models (predict stimulus from EEG) rather than forward models (predict EEG from stimulus), as backward models have been shown to achieve a better fit, at least for envelope-based attention decoding^11^. The mTRF toolbox fits a set of weights using a Tikhonov regularized least squares regression to predict a given stimulus feature from EEG data. To capture the temporal dynamics of the brain’s response to the stimulus feature in question, this EEG signal is lagged in time, resulting in a TRF weight for each channel and timepoint combination in the time-lag window (0-800 ms). We train a separate model for each trial and average over the models of the trials in the training set to obtain the model for a respective crossvalidation fold. For a given training set, the optimal value for the regularization hyperparameter lambda is determined in an inner 3-fold crossvalidation loop from a range of lambda values (*λ* = 2^0^, 2^2^, …, 2^20 [13]^). Note, we modified functions in mTRF toolbox for our analysis to increase training efficiency (e.g., to reduce the number of times the time-lagged variance-covariance EEG matrix *X*^*T*^ *X* is computed).

To evaluate the performance on the test set, a prediction of the attended stimulus feature is generated by multiplying the TRF weights with the observed (time-lagged) EEG. This prediction is then compared to the two features of the candidate stimuli, and the stimulus with the highest (pearson’s) correlation is predicted to belong to the attended stimulus. The trained weights of the backward models reflect the contribution of each timepoint and channel, but cannot be interpreted directly as brain activation in response to the stimulus feature (in the way that the weights of forward models can be^14^). However, Haufe et al. define a way, to transform the backward weights (filter) to an interpretable activation pattern (i.e., TRF), such that the trained models *can* be interpreted. We apply this by multiplying the weights with the covariance of the EEG and the inverse of the covariance of the feature vector *y* (note, this is slightly different from Haufe et al. eq. 6, as in the supervised case the latent factors, *s*, and feature vector, *y*, are equivalent).

### Statistics

To compare the accuracies between two models, we run a (Monte Carlo) randomization test. We use the difference in accuracy between the two models as the test statistic, and obtain a null distribution for this difference by computing this test statistic across data where we shuffle predictions across the two methods. Specifically, we swap the prediction status (i.e., correct/incorrect) of a given trial and subject, between the two methods under consideration, with probability 0.5. We take the data across durations into account by swapping the subtrials (of 15 or 30s) based on whether the full trial (60s) was swapped. We then recompute the accuracy for each method, across all subjects, trials, and durations. We run a total of 1000 of such randomizations, computing the test statistic (difference in accuracy between the two methods) for each, resulting in a null distribution for the hypothesis that the accuracies from the two models are drawn from the same distribution. We compare the observed test-statistic against this null distribution to obtain a p-value (*P*(*t_null_* ≥ *t_obs_*)). To account for the fact that we generate only a subset of all possible randomizations, a more conservative p-value is calculated by including the observed test statistic in the null distribution (*p* = ([*t_null_* ≥ *t_obs_*] + 1)/(*n* + 1)^27, 28^).

To determine statistical significance between weight vectors/TRFs (representing channels by time), we use non-parametric cluster-based permutation tests^19^ (as implemented in Fieldtrip^29^). Such a test allows for the combination of information across electrodes and timepoints to increase sensitivity of the statistical test without having to correct for multiple comparisons with respect to those aspects. These tests were performed using a within-subject design, and a dependent samples t-test as test-statistic. Note that this non-parametric cluster-based permutation test does not rely on assumptions with regard to the distribution of the data, regardless of the chosen test-statistic (the test-statistic is merely used to quantify the difference between datapoints). We performed one-tailed tests (alpha = 0.05), using 1000 permutations, supplying all channels and timepoints as specified. This cluster-based permutation test outputs the clusters it has identified, if any. Each identified cluster has an associated p-value. The test is determined significant when at least one cluster has a p-value below the alpha level. If one or more clusters are found, we report the smallest p-value (whether significant or not).

